# Disease-specific loss of microbial cross-feeding interactions in the human gut

**DOI:** 10.1101/2023.02.17.528570

**Authors:** Vanessa R. Marcelino, Caitlin Welsh, Christian Diener, Emily L. Gulliver, Emily L. Rutten, Remy B. Young, Edward M. Giles, Sean M. Gibbons, Chris Greening, Samuel C. Forster

**Affiliations:** Department of Molecular and Translational Sciences, Monash University, Clayton, VIC 3168, Australia; Centre for Innate Immunity and Infectious Diseases, Hudson Institute of Medical Research, Clayton, VIC 3168, Australia; Department of Microbiology, Biomedicine Discovery Institute, Clayton, VIC 3800, Australia; Institute for Systems Biology, Seattle, WA 98109, USA; Department of Paediatrics, Monash University, Clayton, VIC 3168, Australia; Bioengineering and Genome Sciences Departments, University of Washington, Seattle, WA 98195, USA; eScience Institute, University of Washington, Seattle, WA 98195, USA

## Abstract

Many gut microorganisms critical to human health rely on nutrients produced by each other for survival; however, these cross-feeding interactions are still challenging to quantify and remain poorly characterized. Here we introduce a Metabolite Exchange Score (MES) to quantify those interactions. Using metagenome-wide metabolic models from over 1600 individuals, the MES allowed us to identify and rank metabolic interactions that were significantly affected by a loss of cross-feeding partners in 10 out of 11 diseases. When applied to a Crohn’s disease case-control study, our approach identified a lack of species with the ability to consume hydrogen sulphide as the main distinguishing microbiome feature of disease. We propose that our conceptual framework will help prioritize in-depth analyses, experiments and clinical targets, and that targeting the restoration of microbial cross-feeding interactions is a promising mechanism-informed strategy to reconstruct a healthy gut ecosystem.

## Introduction

The human gut contains hundreds of microbial species forming a complex and interdependent metabolic network. Over half of the metabolites consumed by gut microbes are by-products of microbial metabolism^1^ with the waste of one species serving as nutrients for others^2–4^. Species interdependence can render microorganisms vulnerable to local extinction if a partner is lost^5^ unless alternative species are available to fill that niche. In this context, having functionally redundant species with the ability to produce or consume the same nutrients is beneficial for the host. While it is generally accepted that high functional redundancy is a characteristic of resilient human gut microbiomes^6–8^, the human health impacts of redundancy in metabolic interactions remain largely uncharacterized. Restoring the diversity of cross-feeding microbial partners represents a logical but still largely unexplored rubric to fight a wide range of diseases linked with an unbalanced gut microbiome.

Mechanistic models that simulate microbial metabolism *in silico* hold the promise to fill our knowledge gap on microbial metabolic interactions^4,9^. Genome-scale metabolic models (GEMs) are based on increasingly comprehensive databases linking genes to biochemical and physiological processes^10,11^. These models have been used to estimate metabolic exchanges between pairs of bacterial species for over a decade^12,13^. Developments in automating the reconstruction of GEMs^14^, manually curating GEMs for thousands of gut microorganisms^15,16^, and in the availability of tools to model interactions between multiple species^17^ have paved the way to build metabolic models for complex microbial communities. Studies using community-wide metabolic models have found dozens to hundreds of significantly different metabolic exchanges in the gut microbiome associated with type 2 diabetes^18^ and in inflammatory bowel disease^19^ when compared to healthy controls. A method to rank these metabolic interactions according to an ecology-based framework provides the opportunity to generate targeted hypotheses underlying mechanistic links between the gut microbiome and diseases.

Here, we introduce a metabolite exchange score derived from metagenome-wide metabolic models, designed to identify the potential microbial cross-feeding interactions most affected in disease. We apply our conceptual framework to an integrated dataset of 1,661 publicly available stool metagenomes, encompassing 15 countries and 11 disease phenotypes. Our framework identified both known and novel microbiome-disease associations, including a link between colorectal cancer and the microbial metabolism of ethanol, a connection between rheumatoid arthritis with microbially-derived ribosyl nicotinamide, and links between Crohn’s disease and specific bacteria that metabolise hydrogen sulphide. The scoring system helps quantify and identify context-dependent disruptions of microbial interactions, which may present as targets of microbiome-based medicines.

## Results

### Potential cross-feeding interactions quantification

To understand the link between cross-feeding interactions and disease we designed the Metabolite Exchange Score (MES). MES is the product of the diversity of taxa predicted to consume and taxa predicted to produce a given metabolite, normalized by the total number of involved taxa (Fig. 1a and methods). The potential production, consumption and exchange of metabolites by each microbiome member is estimated through community-wide metabolic modelling. As with a centrality measure of a network that defines their most connected nodes, metabolites with high MESs are likely to be key components in the microbial food chain. At the other extreme, metabolites where the MES is zero are not produced or not consumed by any member of the community. By comparing MESs for each metabolite across healthy and diseased microbiomes, one can rank and identify the metabolites most affected by the loss of cross-feeding partners (Fig. 1b). Once metabolites have been prioritized with MESs, it is then possible to integrate taxa abundances and their estimated metabolic fluxes to retrieve a consortium of species that act as the main producers or consumers of the targeted metabolites. We propose this approach as a hypothesis generation strategy to guide new discoveries, targeted experiments and clinical trials.

**Figure 1.**
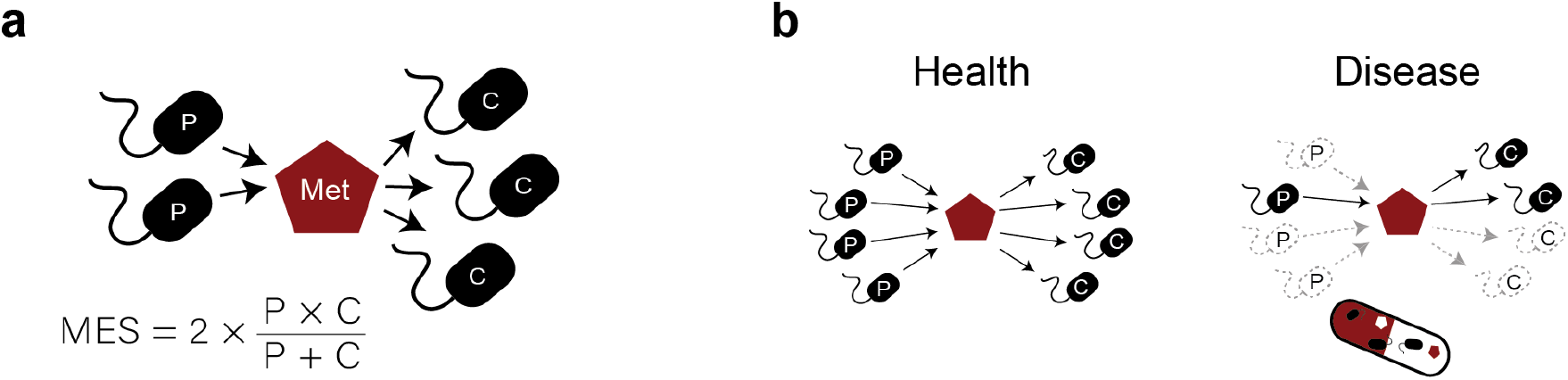
Overview of the Metabolite Exchange Score (MES) calculation and application. **a** The Metabolite Exchange Score is the harmonic mean between the number of potential producers (P) and consumers (C) inferred from metagenome-wide metabolic models. **b** Comparative analysis of MES between healthy and diseased cohorts can help identify the species and metabolites required to restore cross-feeding interactions, which may be promising targets of microbiome therapies.

### Meta-analysis of 1,661 microbiomes reveals key metabolic interactions among gut microorganisms in health and disease

To obtain an overview of the association between cross-feeding interactions and different diseases, we performed a large-scale analysis of 1661 high-quality and deeply sequenced gut metagenome samples, including 871 healthy and 790 diseased individuals from 33 published studies, 15 countries and 11 disease phenotypes (Fig. 2a, Supplementary table S1). Integrating studies and countries enabled the assembly of Metagenome-Assembled Genomes (MAGs) for a diverse range of gut microbes and allowed characterization of the baseline MESs in the healthy population. Our healthy cohort was composed of both males and females with a Body Mass Index (BMI) between 18.5 and 24.9 and no reported disease. Samples for which this information was unclear (e.g. disease controls where health status or BMI was not reported) are not included in our dataset (see Methods for details). Within-sample sequence assembly^20^, metagenome co-binning^21^ and quality control^22^ resulted in 24,369 high-quality MAGs with >90% completeness and <0.05% contamination. We selected one representative MAG per species, defined at 95% Average Nucleotide Identity (ANI), resulting in 949 bacterial and 6 archaeal species, encompassing all dominant microbial phyla found in the gut (Fig. 2b, Supplementary table S2). Presence and abundance of these species was determined by mapping sequence reads against the 955 MAGs. Forty bacterial and one archaeal species were exclusively found in diseased individuals (Supplementary table S3a), while healthy individuals harboured 59 bacterial and one archaeal species that were not observed in any diseased individual (Supplementary table S3b). To infer metabolic exchanges between microbes, we reconstructed Genome-Scale Models (GEMs)^14^ for the 955 MAGs, built community-scale metabolic models for each individual based on the species-level abundances using MICOM^17^, and calculated MESs using custom scripts^23^.

**Figure 2.**
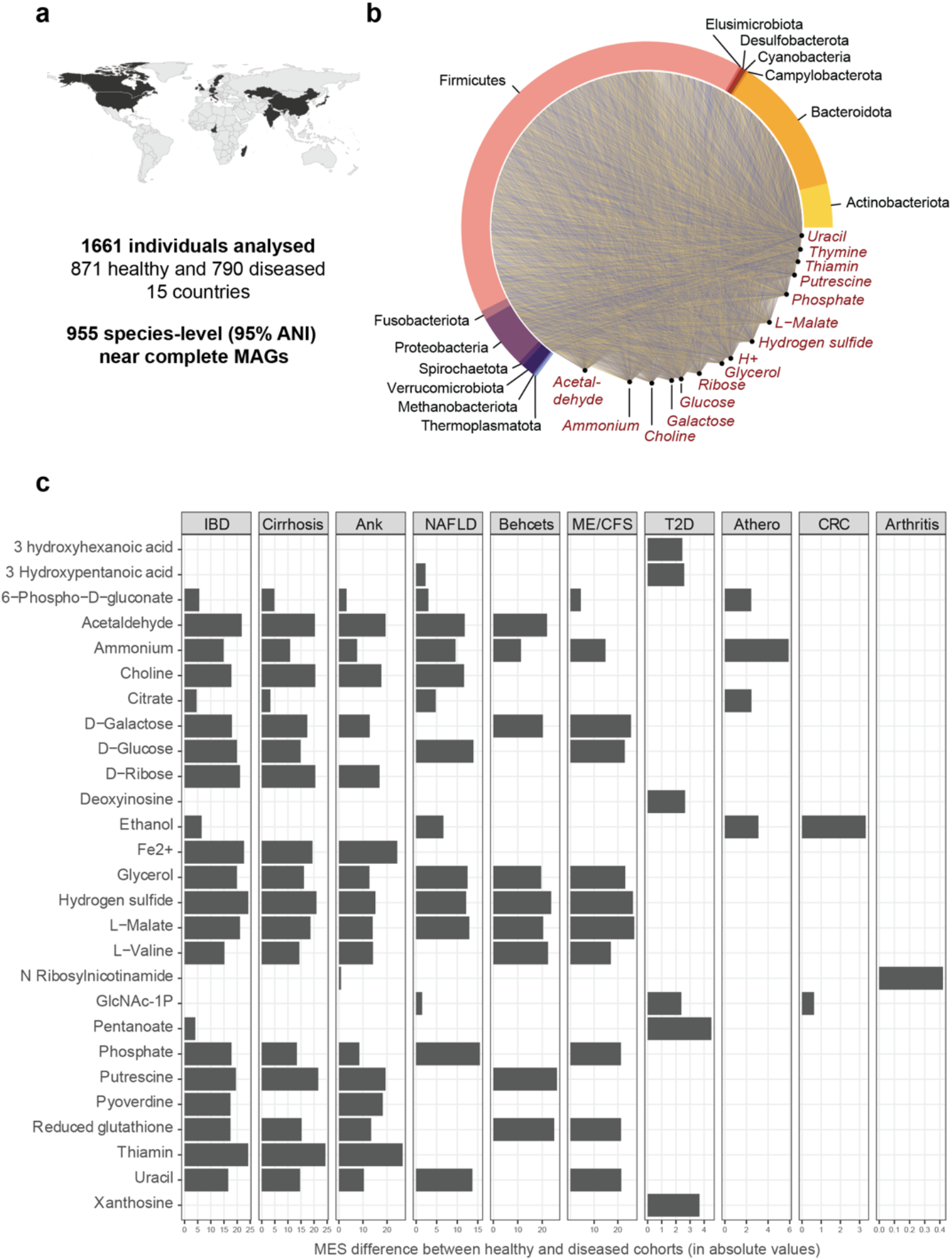
Global analysis reveals most common metabolic exchanges among healthy gut microbes and disease-specific loss of cross-feeding partners. **a** Map highlighting the 15 countries from which metagenomes were included in our analysis. **b** Network of metabolic exchanges within the microbiomes of 871 healthy individuals, highlighting the phyla of the retrieved MAGs and the top 15 metabolites with highest metabolite exchange scores (MES), which are expected to be central to sustain a healthy microbial community structure. **c** Metabolites with significantly reduced MESs in diseased microbiomes when compared to the healthy group (Kruskal-Wallis’ *p* < 0.05/number of comparisons), suggesting significant loss of microbial cross-feeding partners for those metabolites. The panel of metabolites shown here include the top 5 metabolites with highest MES difference between healthy and diseased groups for each disease. No significant difference in MES was found in patients with schizophrenia after accounting for multiple comparisons. Ank=ankylosing spondylitis, IBD=inflammatory bowel disease, NAFLD=non-alcoholic fatty liver disease, ME/CSF=myalgic encephalomyelitis/chronic fatigue syndrome, T2D=type 2 diabetes, Arthero=atherosclerosis, CRC= colorectal cancer, Arthritis=rheumatoid arthritis.

We first sought to identify the metabolic exchanges with the highest diversity of cross-feeding partners in healthy microbiomes by analysing the MESs of the entire healthy group (Fig. 2b). Metabolites showed a wide variation in MES scores between individuals (Supplementary Fig. S1). Metabolites with the highest mean MES included nucleobases such as uracil (MESs mean and sd = 60.5 ± 17.6) and thymine (41.8 ± 21.8), essential nutrients such as phosphate (59.9 ± 17.0) and iron (40.3 ± 36.9), and sugars such as glucose (52.6 ± 22.1) and galactose (52.3 ± 21.3).

To identify the metabolites most affected by the loss of cross-feeding partners during disease, we compared MESs between the healthy group and the eleven disease phenotypes. This analysis identified significant loss of cross-feeding partners for specific metabolites in all disease groups except for Schizophrenia (Fig. 2c, Supplementary Figure S2). Metabolites with high MES in healthy individuals and known to be important for human health, such as vitamin B1 (thiamin)^24^ and precursors of short-chain fatty acids (e.g. malate, glucose, galactose)^25^, were significantly affected in multiple disease phenotypes (Kruskal-Wallis’ *p* < 0.05 / number of tests to correct for multiple comparisons). Thiamin was the metabolite with highest difference in MESs scores between healthy and diseased microbiomes in cirrhosis and ankylosing spondylitis, ranking second in Inflammatory Bowel Disease (IBD) (Fig. 2c). Associations between deficiency of thiamine with cirrhosis and IBD have been previously reported^26–28^, but to our knowledge, this is the first indication of a possible microbial-mediation of this phenotype. Likewise, this is the first indication of a link between microbially-derived ribosyl nicotinamide and rheumatoid arthritis (Fig. 2c). The results also confirmed previously reported microbially-mediated disease-metabolite associations, such as ethanol in colorectal cancer^29^ and hydrogen sulphide in IBD^30,31^, reinforcing the potential of our novel approach to identify reasonable relationships *a priori*.

### Species diversity has distinct relationships with producers and consumers of exchanged metabolites

Diversity of microbial species within the gut community is commonly considered a marker of health status. The number of microbial species exchanging metabolites naturally correlates with the number of species in the community. To further understand the relationship between diversity and metabolite exchange, we tested the null hypothesis that producers and consumers are equally affected by species diversity. Specifically, we correlated the number of producer or consumer species of each metabolite with species diversity to determine statistical differences between the slopes of these correlations for metabolite production and consumption. The null hypothesis (no statistical difference between slopes) implies that the number of producer species and consumer species increases at the same rate as species diversity increases. Such results would imply that cross-feeding interactions dependent only on the number of species present in the community. This null hypothesis was rejected for 79% of metabolites exchanged by the gut microbiome (Fig. 3a, Supplementary table S4), with the slope of the correlation being significantly steeper either for consumers (55% of metabolites) or producers (24% of metabolites). From the metabolites with highest MESs, only producers and consumers of glycerol showed no significant difference in response to species diversity (Fig 3 b–p).

**Figure 3.**
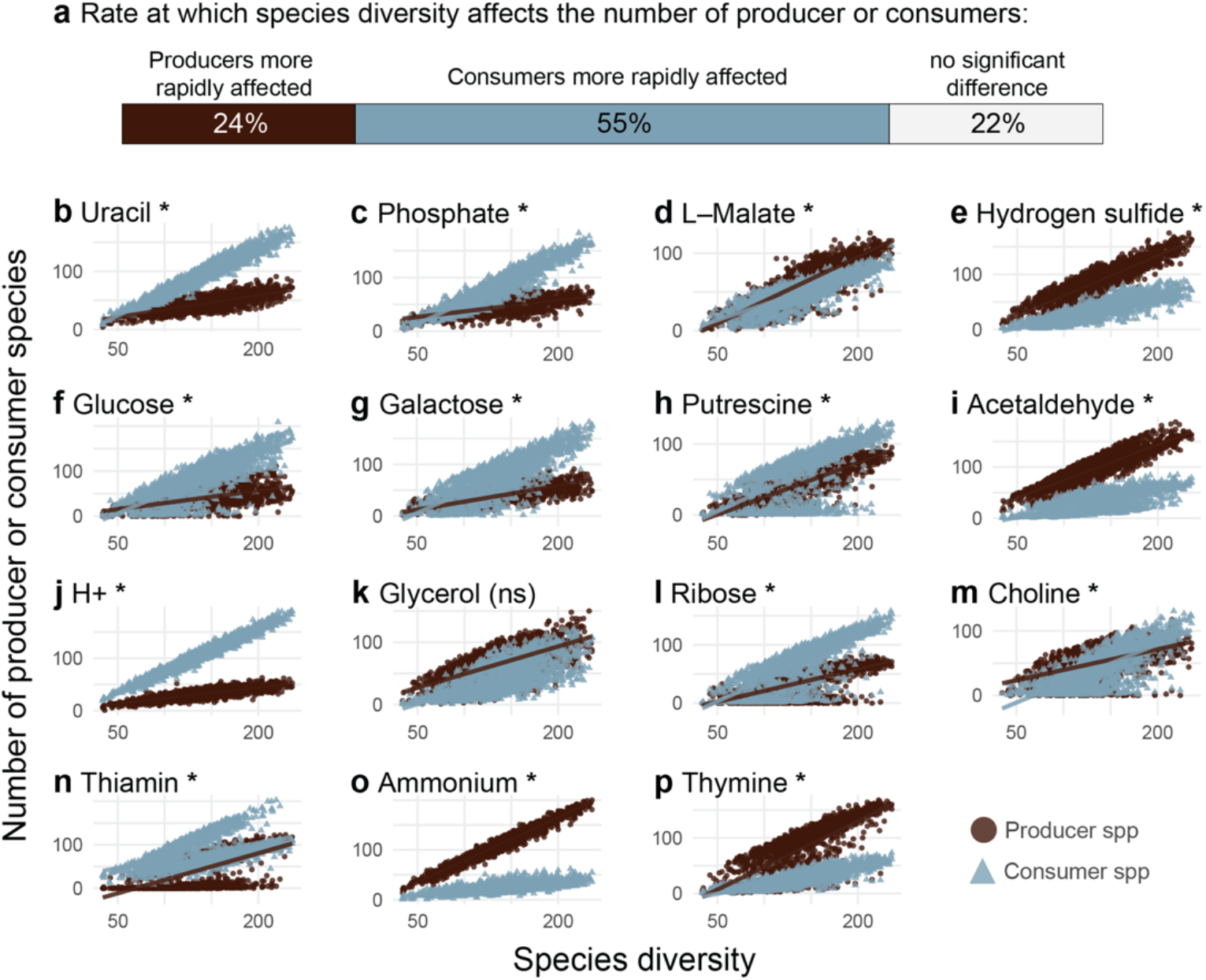
The effects of species diversity on the number of producers and consumers of exchanged metabolic products varies for different metabolites. **a** Differences between the slopes of the species diversity *vs* producers or consumers correlations were observed for the majority of metabolites, with producers having a steeper slope in 24% of the metabolites, and consumers having a steeper slope in 55% of the metabolites analysed. **b-p** Representation of the correlation between species diversity *vs* producers or consumers for the top 15 metabolites with highest MESs in healthy microbiomes. Analyses included all exchanged metabolites present in at least 50 microbiomes (healthy and diseased cohorts). Asterisks indicate a significant *p* value of the regression model after Bonferroni correction.

### Microbial food web restoration as a potential therapeutic strategy for Crohn’s disease

To investigate how the application of Metabolite Exchange Scores and our modelling framework may guide the identification of promising therapeutic targets, we focused on Crohn’s disease (CD), a form of IBD. We selected a single case-control study^32^ with the largest number of samples from healthy and diseased within our quality-controlled dataset to minimize batch effects. In accordance with the global analyses, we found that hydrogen sulphide (H_2_S) – a gas previously implicated in CD and IBD symptoms^30,31,33^– was the metabolite most affected by the loss of cross-feeding microbial partners (twofold reduction, Supplementary table S5). While H_2_S production by the gut microbiome has been subject of several studies (e.g.^34,35^), the consumption if this gas is less characterized, and our modelling results indicate that H_2_S consumed by bacteria can be incorporated into sulphur-containing amino acids such as cysteine (Supplementary Figure S3).

Focusing on H_2_S, we found that the microbiome of healthy individuals contained more species with the potential to produce H_2_S, as well as more species with the potential to consume H_2_S, than the microbiomes associated with CD (Fig. 4a). Interestingly, the diversity of potential H_2_S consumers was more affected in CD patients (56% less diverse on average, supplementary table S6) than the diversity of H_2_S producers (32% less diverse), resulting in a significantly higher H_2_S producer to consumer ratio in individuals affected by CD (Fig. 4c). We observed similar results when investigating the flux of H_2_S among microorganisms. The total estimated ability of the microbiome to consume H_2_S in the disease state was reduced by 74%, while the total production was not significantly affected, resulting in a higher H_2_S production to consumption ratio in CD (Fig. 4b and 4d, Supplementary table S6). The excess of H_2_S (i.e. H_2_S predicted to be exported to medium) was not significantly different between healthy and diseased subjects (Kruskal-Wallis *p* = 0.8). The indication that H_2_S consumers are more affected than H_2_S producers in CD stands after correcting for the confounding effects of species diversity, although no significant difference was observed for the flux of H_2_S exchanged among microorganisms (Supplementary table S7).

**Figure 4.**
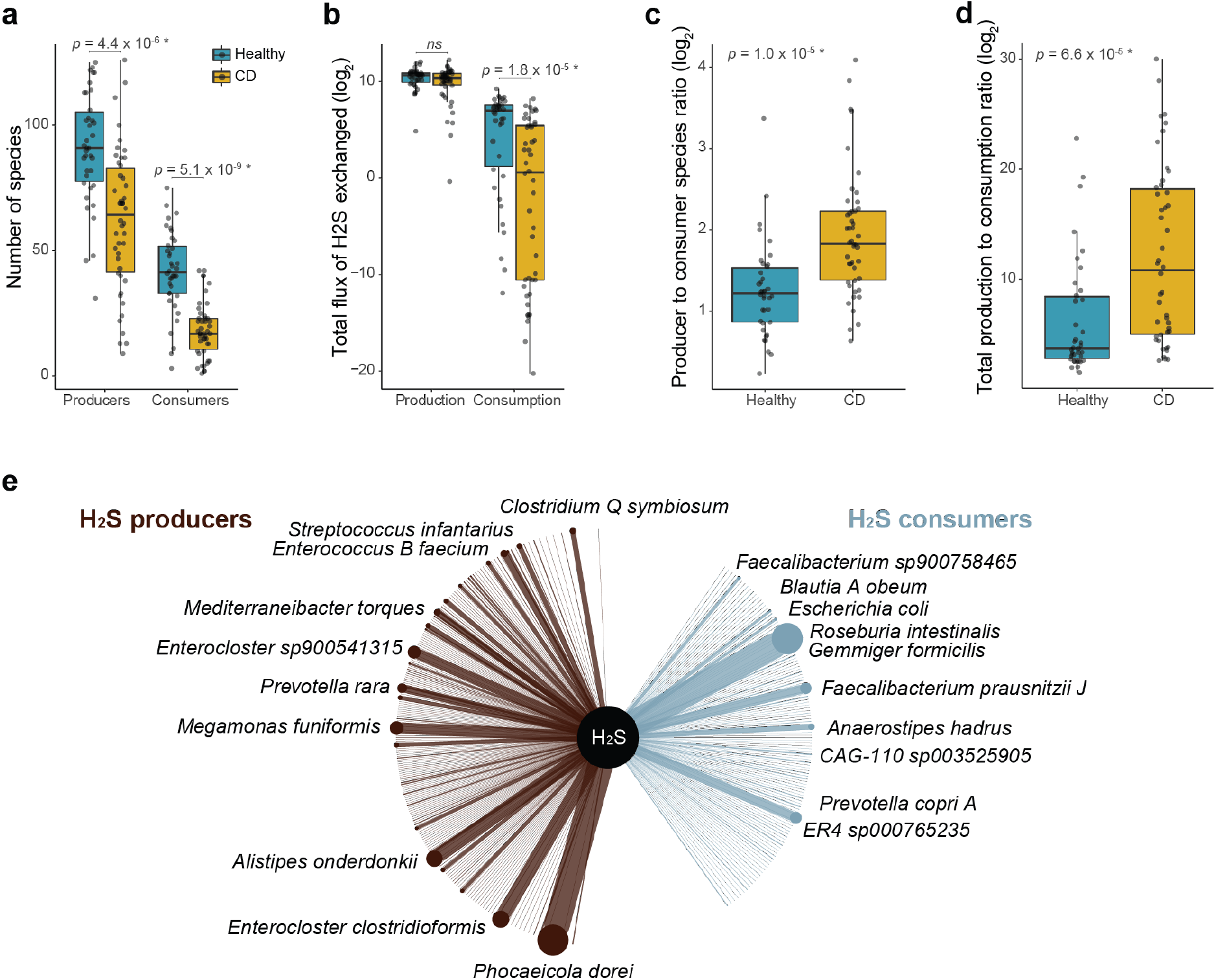
Shift in hydrogen sulphide production-consumption equilibrium associated with Crohn’s disease. **a** The number of species with potential to produce or consume H_2_S is significantly reduced (Kruskal-Wallis *p*-values < 0.05) in microbiomes associated with CD when compared to healthy controls. **b** The total estimated consumption of H_2_S is depleted in CD, while production was not significantly affected (fluxes estimated in millimoles per hour per gram of dry weight). **c-d** A significant increase in the ratio of number of producers to consumers (c) and in the total estimated H_2_S production to consumption (d) was found in microbiomes associated with CD. **e** Species involved in the exchange of H_2_S that are most altered in CD, which might be promising targets of microbiome therapy. The network shows the H_2_S producers with increased production (brown), and the consumers with reduced H_2_S consumption (blue) in CD when compared to healthy controls. The 10 species most affected in each category are highlighted. The thickness of the nodes and edges are proportional to the species’ weighted flux sum of H_2_S within the consumer or producer categories.

To better understand the genetic basis of the metabolic modelling results, we investigated the distribution of 46 genes known to be involved in H_2_S cycling^35^ in the MAGs present in the CD case-control study (Supplementary table S8). We found between one and 23 genes in each MAG (Supplementary table S8). Five genes involved in H_2_S cycling were significantly more prevalent in microbiomes associated with healthy individuals (Supplementary table S9): *cysK, dcm, Fuso_cyst, metH* and *metK* (*p* < 0.0012 accounting for multiple comparisons and using species diversity as a confounder variable). Another five genes were more prevalent in CD-associated microbiomes: *asrA, asrB, asrC, dmsA* and *dsrC* (*p* < 0.0012), the first four genes also being significantly enriched when accounting for species abundance (Supplementary table S9).

To identify the key species associated with H_2_S imbalance in CD, we compared the contribution of each species to the total H_2_S production or consumption in the healthy and CD cohorts. For each species, H_2_S flux (weighted by relative abundances) was estimated and the difference of total H_2_S weighted flux in healthy and CD individuals calculated. The species showing the highest increase towards H_2_S production in CD patients included members of the classes Clostridia, Bacteroidia and Bacilli (Fig. 4e, Supplementary table S10). *Enterocloster clostridioformis* (Clostridia) and *Enterococcus_B faecium* (Bacilli) were only observed in the CD cohort. Many species (45% of the MAGs from the case-control study) showed an ability to both produce and consume H_2_S according to the models, and their role was dependent on their community context. *Phocaeicola dorei* (Bacteroidia) was the species showing the highest difference in predicted H_2_S production between healthy and CD individuals despite being common in both cohorts. We found multiple genes related to H_2_S metabolism in this species (*cysK, bsh, dcm, Fuso cyst, luxS, metK, sufS*, and two copies of the *malY* and *metH* genes). Members of the Clostridia class were the H_2_S consumers showing the highest reduction in H_2_S consumption in CD microbiomes, including *Roseburia intestinalis, Blautia_A obeum*, and two *Faecalibacterium* species (*F. prausnitzii_J* and *F. sp900758465*) (Fig. 4e, Supplementary table S10). The top 5 consumer species had between two and four copies of the cysteine desulfurase (*IscS*) gene, in addition to a range of other genes involved in H_2_S metabolism (Supplementary tables S8 and S10).

## Discussion

In this work we introduce a new MES based conceptual framework and apply it to an integrated dataset of metabolic models for 955 gut species from 1,661 publicly available stool metagenomes, encompassing 15 countries and 11 disease phenotypes. This approach revealed a significant depletion of potential cross-feeding interactions in the microbiomes associated with 10 diseases and identified promising therapeutic targets in a case-control Crohn’s disease study.

We show that our analytical framework identifies both known and novel microbiome-disease associations, providing a cost-efficient and mechanistically grounded strategy to prioritize experiments and guide clinical trials. One example is the link between rheumatoid arthritis and ribosyl nicotinamide (also known as nicotinamide riboside, or NR). This metabolite is one of the main precursors of nicotinamide adenine dinucleotide (NAD+), which has been reported to be significantly reduced in individuals with rheumatoid arthritis^36^. Administration of NR and other NAD+ precursors leads to improved clinical outcomes for rheumatoid arthritis patients^36^ and for a range of other inflammatory, neurodegenerative and cardiovascular diseases^37^. To our knowledge, this is the first reported evidence for a role of microbial NR metabolism in rheumatoid arthritis. We also identified ethanol as the metabolite most affected by loss of cross-feeding in individuals with Colorectal Cancer (CRC). Moderate to heavy alcohol consumption is associated with a 1.17 – 1.44 higher risk of developing CRC^38^ via a process that is at least partially mediated by the microbiome, as gut bacteria metabolise ethanol to produce the carcinogenic acetaldehyde^39^. The capacity to identify these and other coherent metabolite-disease links using exclusively metagenome data is further evidence for the validity and utility of our approach.

Assessing microbiome-disease associations from a functional perspective has been shown to reveal patterns that would not be perceptible in a taxonomy-centric analysis^40^. This phenomenon is supported by our analyses that demonstrates the number of consumers of microbially-derived metabolites tend to respond more quickly to species diversity than the number of producers. It is possible that low species diversity limits the availability of metabolites that can be consumed in the next trophic level, driving microbial consumers to local extinction.

Using CD as a case study, we demonstrated how the metagenome-wide modelling framework can help define mechanistically informed hypotheses for targeted experimental and clinical validation. Our results suggest that CD patients lack microbial community members to support a healthy H_2_S balance. This gas is expected to have a protective effect in the gut when present in small amounts, but it disrupts the mucus layer and may cause inflammation when present in larger quantities^41–44^. Our results corroborate recent findings suggesting that the microbiome of IBD patients is particularly deficient in secreting metabolites containing sulphur^45^, and additionally indicate that H_2_S consumer species are disproportionately lost in CD. Microbial exchanges of H_2_S may affect the host directly through mechanisms such as modulating luminal pH^31^, or indirectly through cascade effects on microbiome composition.

The accuracy of the metagenome-wide modelling framework applied here is limited by the use of automated genome-scale metabolic reconstructions, which represent phenotypes close to manually-curated models^14^ but are naturally unable to predict all organism-specific traits, especially if those rely on genes and pathways that are yet to be characterized. Automated genome-scale models provide an opportunity for a top-down approach, where large scale analyses like the one performed here can guide a range of more refined hypothesis-driven studies, ideally coupled with experimental validation. Additional refinement can be obtained in future studies handling smaller datasets by manual model curation and integration of other omics data^e.g. 46^, and by integrating personalized data on host diet and metabolism^47^.

We expect that metagenome-wide metabolic models, coupled with an assessment of microbial cross-feeding interactions, will help alleviate one of the main barriers in the development of microbiome therapies – prioritizing which species or metabolites to target. By focusing on restoring key aspects of the gut ecology, we may be able to introduce more effective and long-lasting changes in the human gut microbiome.

## Methods

### Global survey of gut metagenomes and quality control

We performed a literature search for peer-reviewed studies with publicly available human stool metagenomes and associated metadata. These included large-scale meta-analyses of gut metagenomes and metadata compilations^48,49^. Studies focusing on dietary interventions, medications, exercise and children (<10 years old) were excluded. For longitudinal studies, only one sample per individual was included in the analyses. To minimize the impact of sequencing technologies, only studies reporting paired-end sequencing using Illumina’s HiSeq or NovaSeq platforms were included.

The healthy cohort included individuals reported as not having any evident disease or adverse symptoms^49^. Samples classified as disease controls and where the health status could not be determined were excluded. To avoid ambiguous health/disease status, samples from individuals with colorectal adenoma (non-cancerous tumour) and impaired glucose tolerance (pre-diabetes) were excluded, and only individuals with a Body Mass Index (BMI) between 18.5 and 24.9 were included in the healthy cohort. Samples with less than 15M PE reads after quality control were excluded to minimize the impact of sequencing depth. A maximum of 100 samples per disease category from each study were used to minimize batch effects and reduce the dataset to a computationally feasible size.

Raw sequence reads were downloaded from NCBI and subject to quality control with TrimGalore v.0.6.6 (Krueger F. http://www.bioinformatics.babraham.ac.uk/projects/trim_galore/) using a minimum length threshold of 80bp and a minimum Phred score of 25. Potential contamination with human sequence reads was removed by mapping the metagenome sequences to the human genome with bowtie v.2.3.5^50^. To minimize the impact of sequence depth, samples were rarefied to 15M fragments (30M PE reads) with seqtk v.1.3 (https://github.com/lh3/seqtk). The quality-controlled dataset contained 1697 samples, which are provided along with their metadata and SRA BioSample identifiers in Supplementary table S1.

### Metagenome assembly and binning

Assembly was performed for individual metagenomes with Megahit v.1.2.9^20^. Metagenome co-binning was performed with Vamb v.3.0.2^21^, dividing the 1697 samples into two batches due to the high computational requirements of using co-abundance information. Completeness and contamination levels of metagenome bins were assessed with CheckM^22^. We retrieved 24,369 bins with > 90% completeness and <0.05% contamination. These bins were dereplicated at 95%ANI using drep v.3.0.0^51^, which selects the ‘best’ representative genome based on multiple quality metrics (completeness, contamination, strain heterogeneity, N50, centrality). De-replication resulted in 955 high-quality, species-level (95% ANI) metagenome-assembled genomes. These MAGs were taxonomically classified with GTDBtk v.1.5.1^52^ and their species abundances across samples were calculated by mapping sequence reads to MAGs with KMA v.1.3.13^53^.

### Genome and metagenome-scale metabolic modelling

Genome-scale metabolic models (GEMs) were reconstructed for each species-level MAG with CarveMe v1.5^14^. GEMs were produced using domain-specific templates for archaea and bacteria, an average European diet^54^ as medium for gap filling, and the IBM Cplex solver.

Metabolic exchanges between community members of a microbiome were calculated with MICOM v.0.26^17^. MICOM simulates growth and metabolic exchanges among members of the microbiome while accounting for their differential abundances, and it has been shown to estimate realistic growth rates. Furthermore, MICOM is computationally tractable when it comes to simulating diverse microbial communities (i.e., dozens-to-hundreds of species). Metabolic exchanges were estimated with MICOM’s growth workflow, using a 0.5 trade-off parameter, an average European diet as medium, and parsimonious Flux Balance Analysis (pFBA) to identify optimal growth rates. The underlying CarveMe models contain relatively few carbon sources, leading to low growth rates and consequent numerical instability. Therefore, the fluxes of medium items were multiplied by 600 to feasibly calculate metabolic exchanges, and then corrected in the final results. An optimal solution was not found for 36 samples, which were removed from the analysis, resulting in a final dataset of 1661 samples. A snakemake workflow is provided in the Zenodo repository for reproducibility^23^.

### Metabolite Exchange Scores

The underlying rationale to define the Metabolite Exchange Score (MES) is that an individual where metabolites are produced and consumed by multiple members of the microbiome will have a higher functional redundancy than an individual where these metabolites are produced and consumed by fewer species, which is a characteristic of most healthy ecosystems. For homogenized stool-derived metagenomes, which do not capture the patchiness in microbial aggregates found in the gut, high functional redundancy increases the likelihood that most micro-niches are populated by at least one species. The MES weighs the number of microbial species consuming and producing a given metabolite, in a given microbiome sample. MES was defined for each metabolite as the harmonic mean between potential consumers and producers:

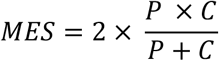

Where *P* is the number of potential producers and *C* is the number of potential consumers of a given metabolite. Note that MES will be zero if a metabolite is only produced or only consumed but not exchanged among microorganisms.

The specific metabolites for which cross-feeding partners were significantly lost were identified with a Kruskal-Wallis test comparing diseased phenotypes against the healthy population. The Bonferroni method was used to account for multiple tests (0.05 as target alpha, divided by the number of tests), and only metabolites present in at least 50 individuals, including at least 15 diseased subjects, were included in the analyses. Water and oxygen were excluded from the analyses. For a simplified graphical representation (Fig 2c), metabolites were selected for display if they showed a significant reduction in the number of cross-feeding partners, and if they were in the top 5 metabolites with highest difference in MES in any disease. Barplots were generated with the *ggplot2* R package^55^. An additional word cloud including up to 100 metabolites with significant MES differences between healthy and diseased was generated with the *wordcloud* R package^56^.

### Species diversity effects

We used the total number of species as a measure of species diversity. Differences in species diversity between healthy and diseased microbiomes were assessed using the Wilcoxon test.

Differences in the slopes between species diversity and consumer or producer correlations were assessed by fitting a linear model in R, considering the interaction between number of producers and consumers with their category (producer or consumer):

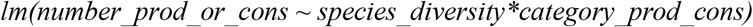

The statistical significance for the difference between slopes was corrected for multiple comparisons using the Bonferroni method.

### Nutritional interactions in the microbiome associated with Crohn’s disease

We selected a case-control study for an in-depth analysis that demonstrates how our framework can be applied to identify promising therapeutic targets. Given that the completeness of metagenome-assembled genomes is optimized by co-binning large datasets^21^, we opted to select a case-control study from our quality-controlled dataset to take advantage of the large number of high-quality MAGs used to model community-wide metabolism. A total of 84 samples from the study of He and colleagues^32^ – the largest CD study within our dataset – passed our quality control and were included in our analyses, including 46 patients with Crohn’s disease and 38 healthy controls. The specific metabolites for which cross-feeding partners were lost were identified with a Kruskal-Wallis test, using only metabolites observed in over half of the samples and adjusting for multiple tests with a Bonferroni correction.

The flux of H_2_S, estimated in millimoles per hour per gram of dry weight, was multiplied by species abundances to obtain the total H_2_S production and consumption exchanged among microorganisms. Fluxes were log_2_-transformed for the statistical tests and graphical representation. Differences between the diversity of H_2_S producers and consumers, ratios of producers to consumers, and their fluxes was evaluated with Kruskal-Wallis tests. The H_2_S predicted to be exported to medium was used to estimate the excess H_2_S production by the microbiome.

We used a nested linear model to account for the confounding effects of species diversity on the associations between number or flux of producers/consumers and disease status. Samples containing less than 99 species (the minimum number of species in the healthy cohort) were excluded from this analysis (n=58 samples remaining), ensuring a linear relationship between species diversity and number of H_2_S consumers or producers.

To better understand the genetic basis of H_2_S production and consumption in MAGs observed within the CD case-control study, we performed a Hidden Markov Model (HMM) survey of 74 genes involved in H_2_S cycling^35^ with HMMer v.3.3.2^57^, using trusted cutoff scores to ensure homology. We used a linear model to test if these genes were differentially distributed between healthy and CD individuals, using only samples with at least 100 species and genes observed in at least 10 samples. Analyses were performed considering both MAGs abundance (by multiplying gene counts by spp. abundance) and prevalence (using species presence/absence, which would be more informative when relatively rare taxa are responsible for a large proportion of the production and consumption of H_2_S). Data was offset by 0.1 to avoid infinity upon log-transformation, species diversity was used as a confounding variable and the Bonferroni correction was used to account for multiple comparisons.

In order to identify species that may be promising targets of microbiome therapy in CD, we weighted in their flux of H_2_S and relative abundances within CD and healthy cohorts. Specifically, weighted H_2_S fluxes of each microbial species was estimated by multiplying their H_2_S fluxes by their relative abundances. The weighted sum of H_2_S fluxes was calculated as the sum of all weighted fluxes within healthy or diseased cohorts. Differences in the weighted sum of H_2_S between healthy and CD cohorts pointed to the key H_2_S producers and consumers associated with Crohn’s disease. The Crohn’s disease cohort contained more individuals than the healthy one, therefore eight random samples were excluded to ensure the same number of individuals (38) in healthy and diseased categories. The metabolic model of *Roseburia intestinalis*, one key H_2_S consumer, was visualized with Fluxer^58^ using best *k-*shortest paths to visualize pathways between H_2_S intake and cell growth.

## Acknowledgments

This work was supported by the Australian Research Council (DP190101504) and the Australian National Health and Medical Research Council (APP1181105 and APP1186371). V.R.M. is supported by an Australian Research Council DECRA Fellowship (DE220100965) and S.C.F. is supported by a CSL Centenary Fellowship. The authors also acknowledge the Victorian government infrastructure support fund and thank the Monash eResearch facility for providing computational resources. We also thank Paul Harrison and Jamie Gearing for statistical and bioinformatics advice.

