## Supplementary material for "Disease-specific loss of microbial cross-feeding interactions in the human gut": Figures S1 - S3

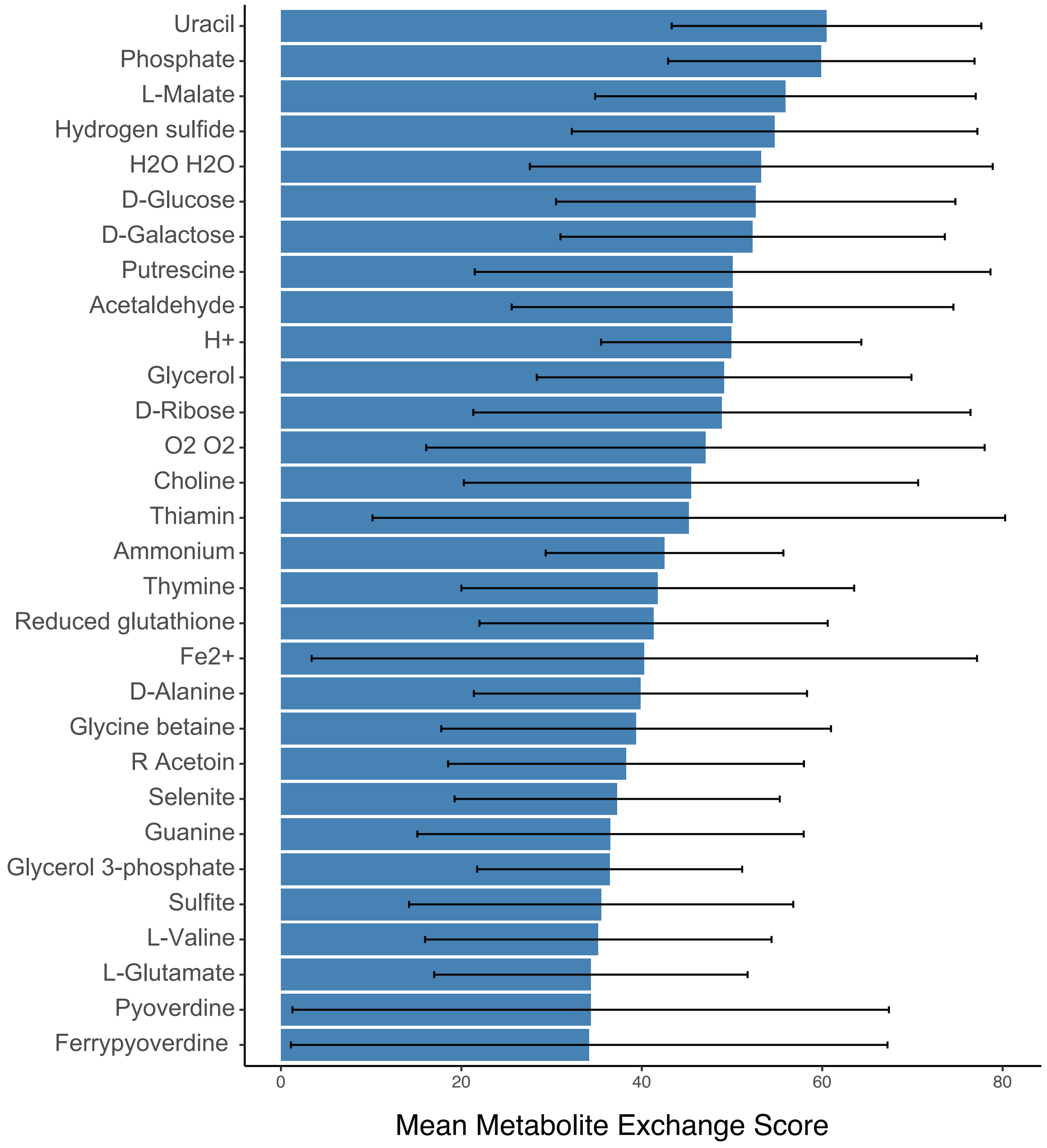


**Supplementary Figure S1.** Top 30 metabolites with highest Metabolite Exchange Score in the healthy human gut microbiome (871 microbiomes).

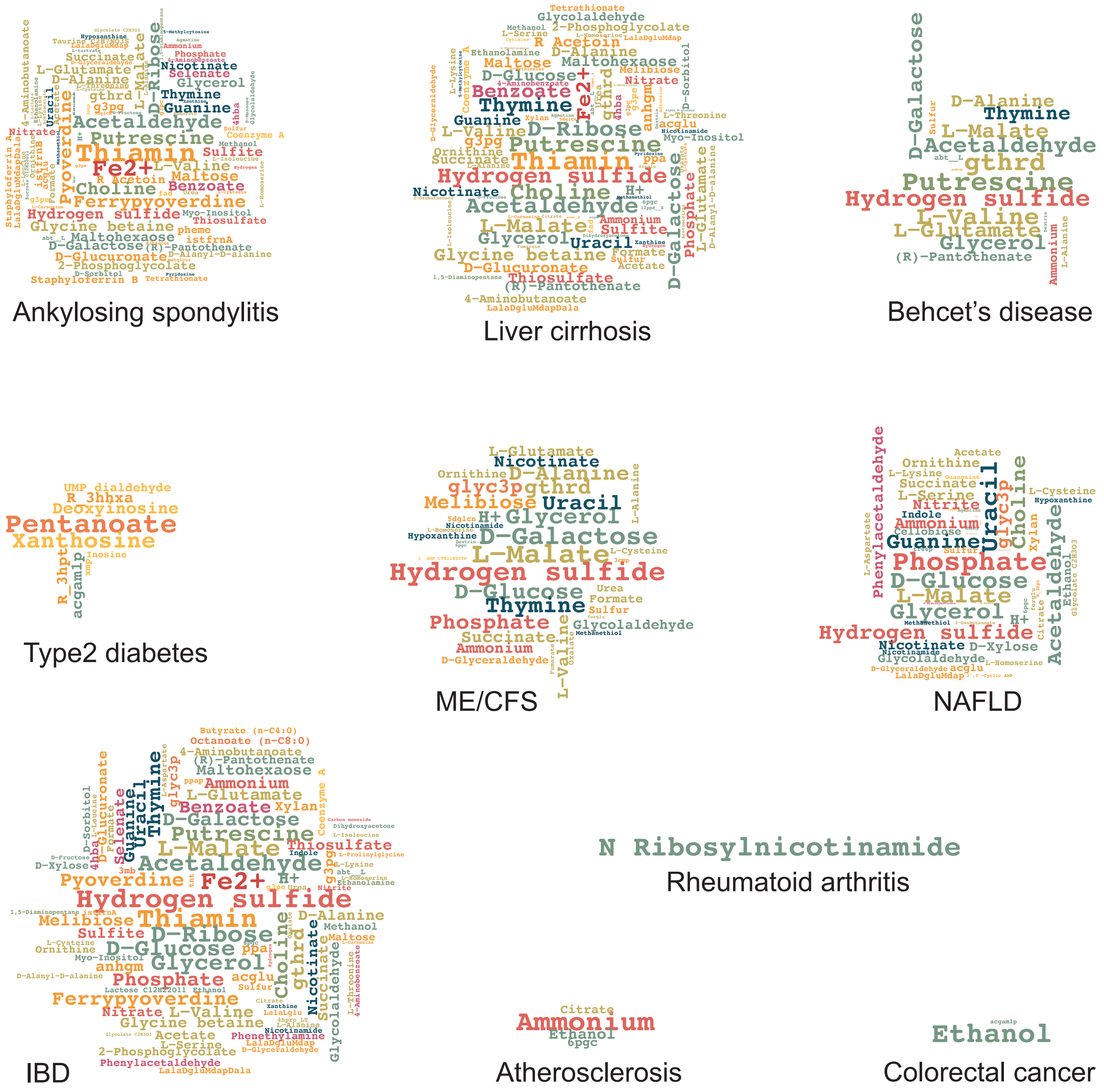


**Supplementary figure S2.** Metabolites with significantly reduced Metabolite Exchange Scores (Kruskal Wallis' p < 0.05/number of comparisons) in the microbiomes associated with 10 disease phenotypes when compared to the healthy group, suggesting significant loss of microbial cross-feeding partners for those metabolites. Letters in word clouds are proportional to the difference in MES between health and disease. While figure 2c (main text) included only the top 5 metabolites with highest MES difference betwee healthy and diseased groups for each disease (for readability), these word clouds include up to 100 metabolites.

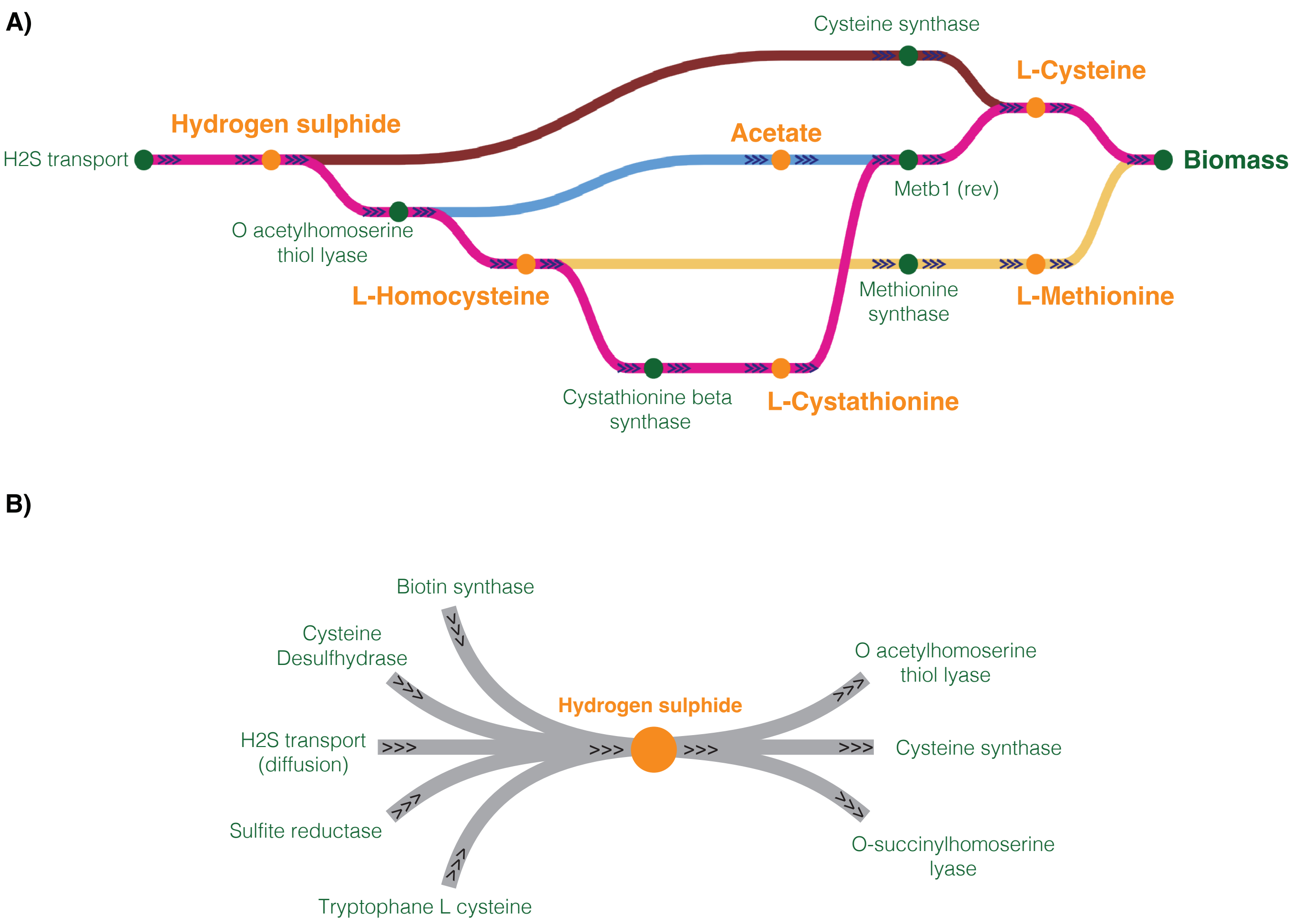


**Supplementary figure S3.** Overview of the metabolic reactions utilising imported hydrogen sulphide in *Roseburia intestinalis*, one of the key $H_2S$ consumers that is depleted in the microbiome associated with Crohn's disease. A) Shortest *k* paths using $H_2S$ transport as source and biomass reaction as target. B) All potential reactions leading to production and consumption of $H_2S$.
